# Cooperation of the *haves* and the *have-nots*

**DOI:** 10.1101/234849

**Authors:** Kaumudi H Prabhakara, Albert J Bae, Eberhard Bodenschatz

## Abstract

Upon starvation, Dictyostelium discoideum (D.d.) exhibit social behavior mediated by the chemical messenger cyclic adenosine monophosphate (cAMP). Large scale cAMP waves synchronize the population of starving cells and enable them to aggregate and form a multi-cellular organism. Here, we explore the effect of cell-to-cell variability in the production of cAMP on aggregation. We create a mixture of extreme cell-to-cell variability by mixing a few cells that produce cAMP *(haves*) with a majority of mutants that cannot produce cAMP (*have-nots*). Surprisingly, such mixtures aggregate, although each population on its own cannot aggregate. We show that (1) a lack of divalent ions kills the *haves* at low densities and (2) the *have-nots* supply the cAMP degrading enzyme, phosphodiesterase, which, in the presence of divalent ions, enables the mixture to aggregate. Our results suggest that a range of degradation rates induces optimal aggregation. The *haves* and the *have-nots* cooperate by sharing complementary resources.

## INTRODUCTION

*Dictyostelium discoideum (D.d.)* is a soil-dwelling amoeba, about 10μm in size with a unique survival strategy. In nutrient rich environments, the cells feed, divide and lead a solitary life, but when food becomes scarce and the cells start to starve, they exhibit social behavior [1, 2]. *D.d.* signal each other by generating excitable waves of the chemical messenger cyclic adenosine monophosphate (cAMP)—when cells at one location sense that the external cAMP concentration is rising due to secretions from their neighbors to one side, these cells are in turn stimulated to secrete cAMP and relay the signal to their neighbors to the other side. To bring the concentrations back down after a wave passes, there is global degradation of cAMP by a phosphodiesterase (PDE) that is continuously secreted by the cells [3, 4]. Excitation centers emerge, and after about 6 hours of starvation, the cells become chemotactic, and migrate towards these centers. Approximately 10^5^ cells come together to form a multi-cellular organism which culminates into a fruiting body composed of a ball of spores atop a stalk. When food is restored, the spores germinate into amoebae. This unique ability of *D.d.* to transform from solitary cells to a multi-cellular organism has made it a much studied model system for morphogenesis [5-8].

Essential to the survival of *D.d.* is its ability to self-organize by chemical signaling. However, in a population, there is significant cell-to-cell variability in cAMP production [9-11]. How then is the signaling mechanism so robust? We intensified the cell-to-cell variability by mixing two kinds of cells, those that can produce cAMP, which we call the *haves,* and the *have-nots,* which are mutants that cannot produce cAMP and thus cannot aggregate. Puzzlingly, when we add a small number of *haves* to a high density of *have-nots,* the mixture aggregates, whereas neither population aggregates on its own. In this paper we solve this puzzle.

## RESULTS

Development of *D.d.* under starvation is typically studied at densities of ≈ 10^5^ cells/cm^2^ on agar [6, 12]. In our experiments, we worked with similar densities of cells, albeit on a Petri dish under a layer of buffer solution instead of agar, because this gave us the greatest uniformity in cell plating density. At these densities, cells initially form a confluent layer on the surface and later aggregate under starvation. When we decreased the density below ≈ 10^4^ cells/cm^2^, the population no longer aggregated. However, when we added about 10^5^ cells/cm^2^ of the acaA^-^ mutants (*have-nots*) to these low densities of wild-type cells (*haves*) aggregation was rescued (figure 1). This is surprising because a pure population of 10^5^ cells/cm^2^ of *have-nots* didn’t aggregate, as they were unable to produce cAMP. Since neither the low-density population of *haves* nor the high-density of *have-nots* was capable of aggregating on their own, there must be a synergy between the two populations that rescues aggregation.

**FIG. 1.**
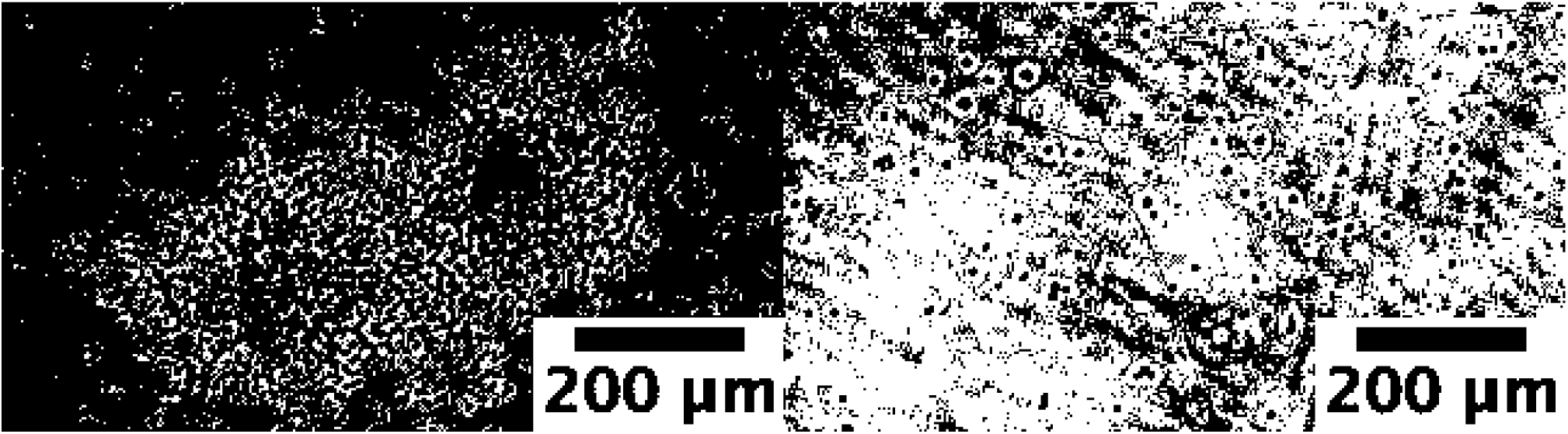
(a) A mixture of the high-density *have-nots* and a low-density *haves* can aggregate, whereas (b) a low-density population of *haves* cannot aggregate on its own. Images were taken about 12 h after start of starvation. These experiments were repeated seven times.

How do these *haves* and *have-nots* cooperate? The cooperation between the two populations can only be chemical or mechanical in nature. Let us first hypothesize that the presence of *have-nots* is sensed by *haves* through mechanical interactions because the *have-nots* don’t produce cAMP, and therefore shouldn’t directly affect the cAMP relay. This mechanical hypothesis is supported by recent work showing that mechanical stresses play a role in signal transduction [13, 14]. To first approximation, we replaced the mutants with 10μm polystyrene beads. If the interactions were due to simple mechanical contact, beads might play the role of the *have-nots.* This, however, we did not observe (in two experiments). The beads did not rescue aggregation. As a positive control, to ensure the beads are not harmful to the cells, we added the beads to a high density of *haves,* where we know the cells do aggregate. We found that the cells not only aggregated, but also actively transported the beads as they moved and incorporated them into their aggregates (figure 2, Supplementary movie S1).

**FIG. 2.**
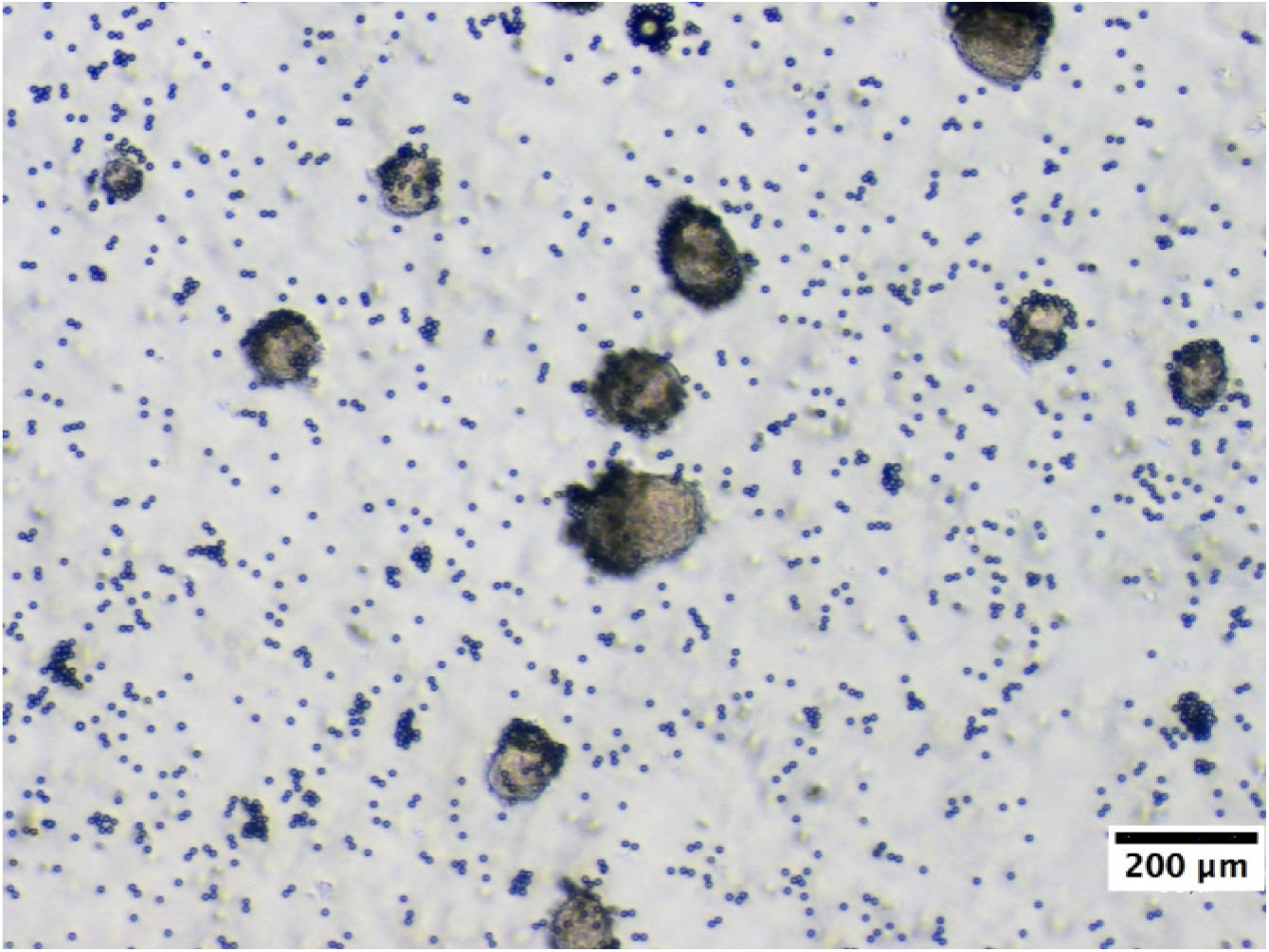
Beads added to a high-density population of *haves* are incorporated into the aggregates. These experiments were performed about 35 times with different ratios of the beads to the high-density *haves.* The beads did not affect pattern formation in all cases.

Other subtler contact-dependent interactions could be in play. To completely rule these out, we placed a Millipore filter between the populations of the *haves* and the *have-nots.* This filter allows the exchange of chemicals, while preventing the cells from coming into mechanical contact. The populations aggregated (in two trials of the experiments). Thus, we have disproved our mechanical hypothesis. The effect must be chemical in origin.

What is this chemical interaction? Could the aggregation be caused by some yet-to-be-identified chemical factors secreted by the high-density cell populations into the buffer? We obtained supernatants from three sources: a high-density population (2 × 10^5^ cells/cm^2^) of *haves,* a high-density population of *have-nots* (2 × 10^5^ cells/cm^2^), and the mixture described above (9 × 10^3^ cells/cm^2^ of wild type cells and 2 × 10^5^ cells/cm^2^ of acaA- mutants). We starved these populations for six hours and then carefully collected the supernatants. Each of these supernatants was added to a low-density population of *haves*. Aggregation was rescued in all three cases (in three repetitions of the experiment), showing that the factors responsible for aggregation are secreted by both *haves* and *have-nots*.

What in the supernatants causes the aggregation? To narrow down the list of chemical factors that could potentially be responsible, we separated the supernatant into two fractions, one containing chemicals > *30kDa* and a fraction < *30kDa,* containing small proteins, ions, and other low molecular weight chemicals (see Methods). We denote these fractions as the High Molecular Weight Fraction (HMWF) and Low Molecular Weight Fraction (LMWF) respectively. To further fractionate the HMWF fraction, we subjected a part of it to heat treatment to deactivate heat sensitive enzymes, and left the rest untreated.

Low densities of *haves* were developed in these fractions. After about 20h, we found (in one experiment):

- Cells + LMWF: The low-density *haves* were viable and appeared polarized, but no tight aggregates were observed. (figure 3)
- Cells + heat-treated, HMWF: The low-density *haves* rounded up and did not appear viable. Furthermore, many cells had de-adhered from the substrate (figure 3).
- Cells + non-heat-treated, HMWF: The low-density *haves* also rounded up and did not appear viable (figure 3).
- Cells + LMWF + heat-treated, HMWF: In this positive control, the low-density *haves* looked polarized (figure 4a).
- Cells + LMWF + non-heat-treated, HMWF: In this second positive control, the low-density *haves* showed clear large-scale streaming (figure 4b).

**FIG. 3.**
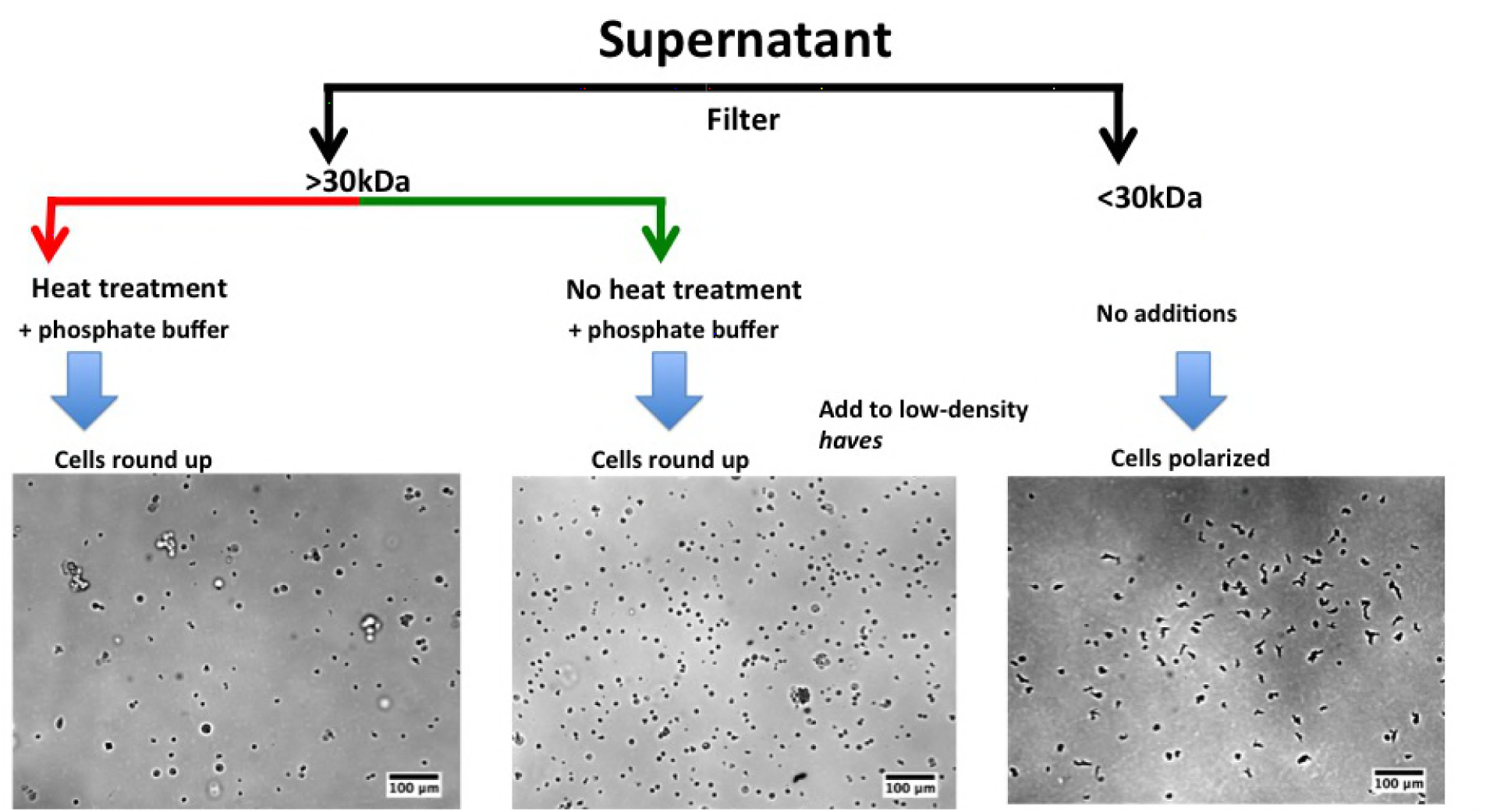
Supplementation of the low-density *haves* with different components of the supernatant. Adding just the HMWF of the supernatant, both heat-treated and non-heat treated, caused the cells to round up. When the LMWF of the supernatant was added, the cells were healthy and polarized.

**FIG. 4.**
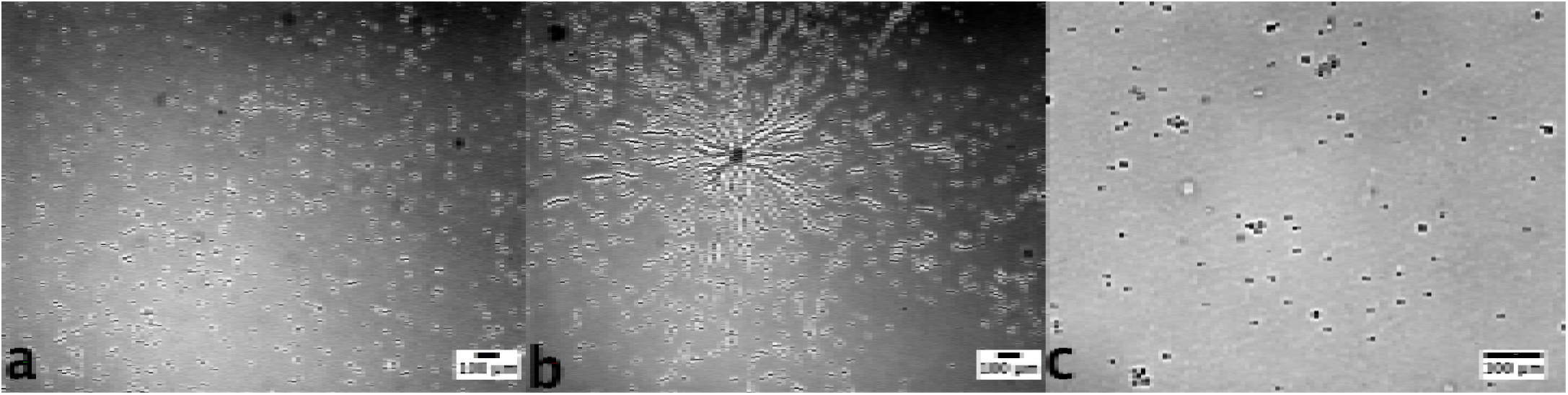
(a) Positive control - heat-treated, HMWF of the supernatant mixed with the LMWF was used to condition the low-density *haves.* (b) Positive control – non-heat treated, HMWF of the supernatant mixed with the LMWF was used to condition the low-density *haves.* (c) Negative control - buffer was added to the low-density *haves.*

We also conducted a negative control, in which the low-density *haves* were unconditioned. These cells rounded up and did not appear viable (figure 4c). Clearly, something smaller than 30kDa is essential to maintain viability. This chemical is secreted by both *haves* and *have-nots,* and is deficient in our buffer. Additionally, a heat-sensitive factor larger than 30kDa enables the cells to signal effectively over long distances.

What could be in the LMWF that keeps the cells alive? We hypothesize that cells carry over essential ions into our low ionic strength buffer. At low densities the ionic strength might not be sufficient for the cells to remain viable [15-17]. To test this hypothesis, we supplemented our buffer with either 50*μ*M CaCl_2_, 50*μ*M MgCl_2_ or 150*μ*M NaCl. For the latter, we used a higher concentration to match the ionic strengths of the other two buffers. In the calcium and magnesium enriched buffers, (figure 5a,b) after 20 h, the low-density *haves* were viable and polarized, but did not form clear aggregates. In contrast, the low-density *haves* in the sodium enriched buffer (figure 5c) rounded up, indicating that they were not viable. We can conclude that divalent ions, Ca^2+^ or Mg^2+^, which are secreted by *have-nots* as well, are required to keep the cells viable.

**FIG. 5.**
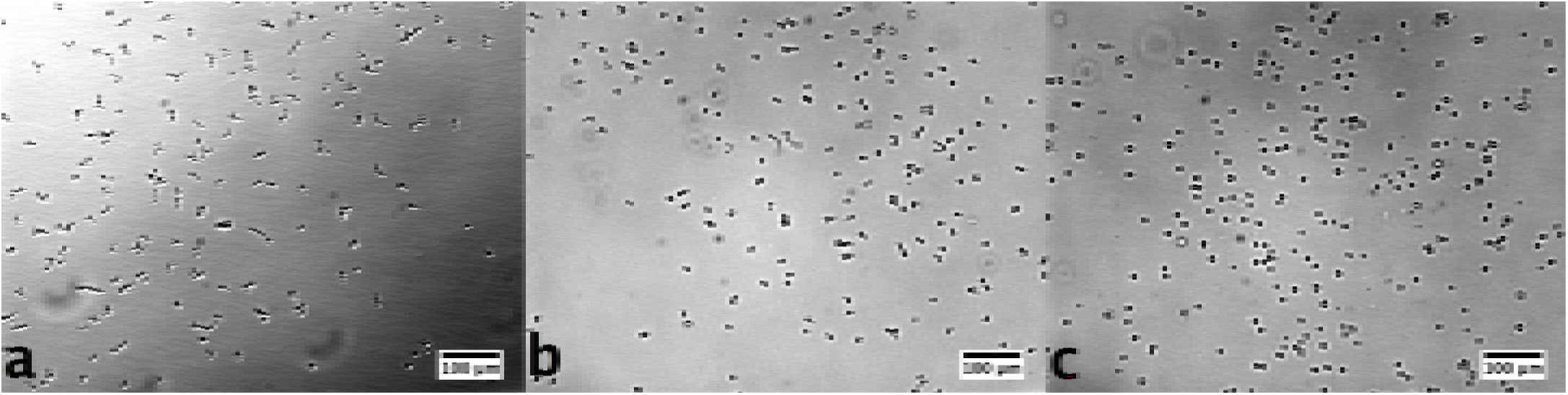
Low density *haves* in (a) calcium enriched buffer were polarized and beginning to stream, (b) magnesium enriched buffer were polarized and beginning to stream, (c) sodium enriched buffer rounded up and were therefore unhealthy. Images were taken a day after start of starvation.The experiments with calcium were performed five times, while the experiments with sodium and magnesium were performed once.

Although the addition of calcium and magnesium kept the cells viable, it did not fully rescue aggregation. This is consistent with the experiment where we added the LMWF of the supernatant to the low-density *haves.* Above, we showed that factors in the heat-sensitive HMWF were necessary to rescue aggregation (figure 4a,b). In the HMWF, we expect to find Conditioned Medium Factor (CMF) and PDE, both of which were shown to be important for aggregation [18-21]. PDE is heat-sensitive [22-24] (see supplementary note S1 for the experimental verification), whereas CMF is not [18]. Could PDE be the missing factor? To demonstrate that this is indeed the case, we added exogenous PDE (PDE4A) at various concentrations to the low-density *haves* conditioned by calcium enriched buffer. We found that at concentrations of PDE above 2 units/ml (see Methods), normal aggregation behavior was recovered (figure 6).

**FIG. 6.**
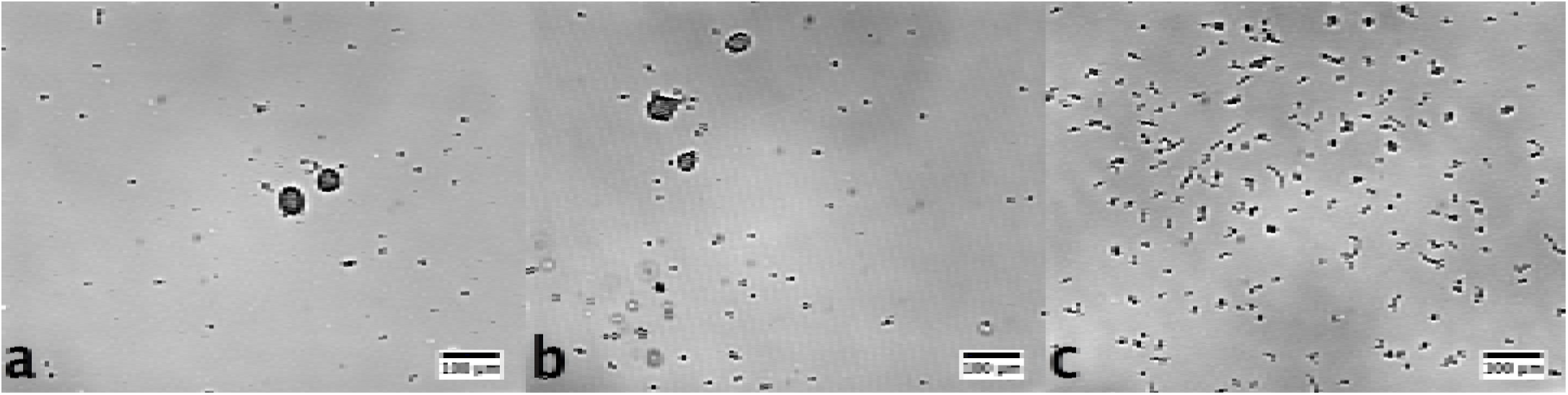
Addition of PDE in different amounts to calcium enriched buffers to condition the low density WT cell populations. (a) 49 units/ml (b) 2 units/ml (c) 0.1 units/ml. Images were taken a day after start of starvation. Experiments were performed once.

## DISCUSSION

We opened this manuscript with a riddle: when a low-density population of *haves* were added to a high-density population of *have-nots*, the mixture aggregated, whereas each population on its own fails to develop.

We found that the lack of divalent ions kills the low-density *haves.* Ions like Ca^2+^ and Mg^2+^ are necessary to keep the cells viable, but they alone do not enable the low-density *haves* to form tight aggregates. Addition of external PDE is necessary for these low-density *haves* to aggregate. Although addition of calcium does induce the cells to secrete more PDE (see Supplementary note S1 and [25-28]), the increased PDE activity due to the addition of calcium is not sufficient to fully restore aggregation in a population of low-density *haves.*

We found that the addition of external PDE to the low-density *haves* in buffers enriched with divalent ions, enabled them to form tight aggregates. Previously, it was shown that pdsA" cells, in which the gene responsible of PDE production has been knocked out, displayed decreased PDE activity and aberrant development and chemotaxis [29-31].

The importance of PDE can be understood by considering wave propagation in populations of *D.d.* The propagation of more than one wave-front requires degradation of cAMP within one wave period, i.e., within about 6-8 min, most of the cAMP should be degraded. As an estimate, if the cells communicate over a length scale corresponding to about 10 cell lengths, which is about 130*μm* at high cell densities, the required degradation rate is about 1.4/ min, which corresponds to 0.7 min. (See Appendix A1 for more details.) Being much smaller than the wave period, this degradation time allows wave propagation, which we observe in the mixture of *haves* and *have-nots* (see supplementary movie S2). If we assume that degradation rate scales linearly with density, then at the low cell density we consider, the degradation rate is about 50 min. This is much longer than the wave period. Therefore well-developed spirals cannot exist, which agrees with what we observe in the low-density population of *haves* in a calcium enriched buffer (see supplementary movie S3).

Addition of exogenous PDE supplements the degradation of cAMP. In our experiments with PDE4A, we estimate the degradation time due to the exogenous PDE to range from 0.2 min to 130 min. (See Appendix A2 for details.) In accordance with our requirements, we observe tight aggregates only in cases where the degradation rate is smaller than or comparable to the wave period, as seen in figure 6.

Having gained these insights, we return to the cooperation between the *haves* and the *have-nots.* The *have-nots* supply divalent ions and PDE (see supplementary note S1), both of which are vital for aggregation. In fact, in a mixture of a 200 cells/cm^2^ of *haves* and 1 × 10^6^ cells/cm^2^ of *have-nots,* so much PDE is produced that waves cannot propagate for long distances - cAMP is completely degraded at short distances (see supplementary movie S4 and figure 7), resulting in small aggregates, where, due to the absence of large scale waves, many cells were excluded. Thus, we see that the degradation rate is an important parameter for aggregation - both at very low and very high degradation rates, aggregation is not efficient.

**FIG. 7.**
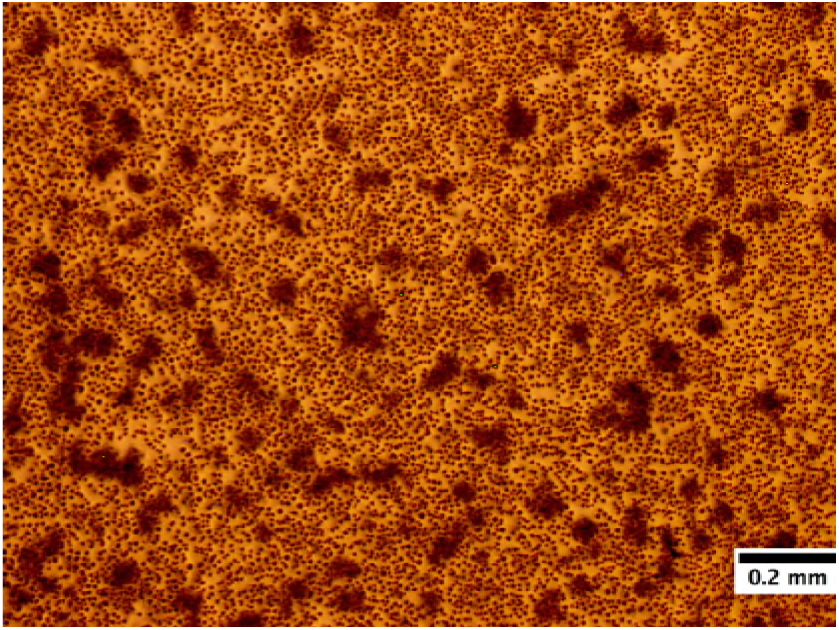
The formation of aggregates in a mixture of a low-density (200 cells/cm^2^) of *haves* and a very high density (1 × 10^6^ cells/cm^2^) of *have-nots.* The *haves* act as centers of activity and the *have-nots* around them aggregated. Due to the high degradation of cAMP by the PDE produced by the *have-nots,* the cAMP waves did not reach all the cells, due to which many cells were not included in any aggregate. This experiment was performed twice. Experiments with similar extreme ratios were performed about 12 times with similar results.

We started this paper by naïvely called the wild-type cells the *haves* and the acaA^-^ cells the *have-nots,* simply because the latter group is incapable of producing cAMP. But we found that the mutants provide other components that are vital for survival. While investigating their cooperation, we found that (1) a critical amount of divalent ions is necessary to keep the cells viable, and (2) PDE activity within a certain range is necessary for optimal aggregation. When these two criteria are satisfied, even a small fraction of cAMP producing cells can synchronize the population and cause aggregation, explaining the robustness of the mechanism. Cooperation between the cells occurs by the sharing of complementary resources. It would be interesting to see whether other species display such cooperation.

## METHODS

### Preparation of the cells

AX2 -214 cells and acaA^-^ mutants (in AX3 background) were grown as Petri dish cultures in HL5 medium (Formedium) at 22°C. These cell lines were kind gifts from G. Gerisch and A. Kortholt respectivelt. The cells were harvested when they became confluent and were washed three times in phosphate buffer (2g of KH_2_PO_4_, 0.36g of Na_2_HPO_4_.2H_2_O per liter at pH 6, autoclaved). The cells were then counted using a hemocytometer and diluted to the appropriate density with fresh phosphate buffer. They were then plated on a 8.6 cm diameter plastic Petri dish with 10 ml phosphate buffer - this corresponds to a buffer thickness of 2 mm. To conserve reagents, we sometimes used plastic Petri dishes divided into quadrants, or smaller diameter Petri dishes. In all these cases, we maintained a constant areal and volume density of the cells. Parafilm was wrapped around the dish lids to prevent evaporation of the buffer. To prevent condensation, we blew 24°C air above the dish lid. Development occurred in a dark room at 22^°^C.

### Supernatants

The supernatant was aspirated from the Petri dishes, and then centrifuged to pellet out any cells accidentally. The supernatant was then cleared using 0.2*μ*M filters.

To fractionate the supernatant into the high and low molecular weight fractions, we used a 30kDa Amicon filter (EMD Millipore). The supernatant was added to the filter and spun down at it 3500 g for about 15 min. This left about 200*μ*L of high molecular weight retentate. The efflux contains the rest of the supernatant which has substances smaller than 30kDa. To use the retentate as conditioning medium at the same concentration as in the supernatant, we reconstituted it in phosphate buffer of the same volume as the starting volume of the supernatant used.

The heat treatment consists of heating the supernatant uniformly at 80°C for about 20 min in a thermostat. The actual temperature was monitored using a thermometer. After 20 min, the supernatant was allowed to cool to room temperature before being used for experiments.

### Addition of external PDE

PDE4A were purchased from Enzo and Signal Chem. For each experiment, the enzyme was diluted based on the activity provided on sample and from the experimental value of the Michaelis-Menten constant for these enzymes. 1 unit = 1 nmol/min at 37°C. For details of the calculations, see Appendix A2.

## ACKNOWLEDGEMENTS

We are grateful to Maren S Müller and Katharina Gunkel for their invaluable support in the preparation of the cells and in procuring the chemicals and enzymes used in this study. We thank Bastian Hüllsmann and the Gorlich lab at the Max-Planck Institute for Biophysical Chemistry for their help and support in the use of their luminometer. We thank J. P. Sethna, V. Vogt, I. Guido and G. Gerisch for useful discussions. We thank C. Westendorf for help with the microscopes and discussions. This work has been supported by the Max Planck Society.

## SUPPLEMENTARY NOTE S1: MEASURING THE EXTRA-CELLULAR PDE ACTIVITY

To measure PDE activity, we used the PDELight kit from Lonza. The measurement works on the principle that PDE degrades cAMP to create AMP. The detection reagent provided in the kit converts AMP to ATP, Luciferin and oxygen. In the presence of luciferase, these combine to produce AMP and photons (among other products). The number of photons measured is thus an indicator of the amount of AMP present. We allowed the PDE from our cell populations to react with cAMP for 20 min and measured the resulting amount of AMP by counting photons with a luminometer. From the amount of AMP remaining, we estimated the degradation of the PDE in our sample.

We first collected the supernatant from the populations as described in Methods. We diluted the retentate of the filtration appropriately. For each sample, we created multiple dilutions of the retentate and added 30*μ*L of each in a black 96 well plate. To each dilution, we added 10*μ*L of 100*μ*M cAMP. After 20 min of reaction time, we added the stop solution - a 1:1 mixture of the Lonza stop solution and 100 mM DTT. Then, we added the detection reagent, and allowed 10 min for the reaction to stabilize. For each dilution, we created a background sample by adding the just diluted retentates without cAMP to the wells in the 96 well plate. During each experiment, for calibration, we added 30*μL* of different concentrations of AMP to the same 96 well plate. To make the same volume, instead of cAMP, we added 10*μ*L of phosphate buffer to these standards. We added the same amounts of stop solution and detection reagent. An integration time of 0.1s was used. We recorded the luminosity every 10 min to account for the decay of AMP. We fit the time series of the intensities with a model (Appendix A3) to obtain the degradation rates.

We performed this assay to first find the degradation rate of cAMP in different cell populations. At high wild type cell densities, the degradation rate is about 0.007 /min (figure 8 column a). At half this density, the degradation rate decreases to 0.0018 /min (figure 8 column b) and at the low density, it is 0.0002 /min (figure 8 column c). The mutants also produce a significant amount of PDE, although it is not as much as that produced by the same number of wild type cells (compare figure 8 columns a and d). Further, keeping the total cell density fixed, as the ratio of mutants to wild type cells is decreased, the population mixture produces more PDE (figure 8 columns d-f). From these measurements it was clear that the mutants contribute PDE in addition to the divalent ions in the mixture of low density wild types and high density mutants.

**FIG. 8.**
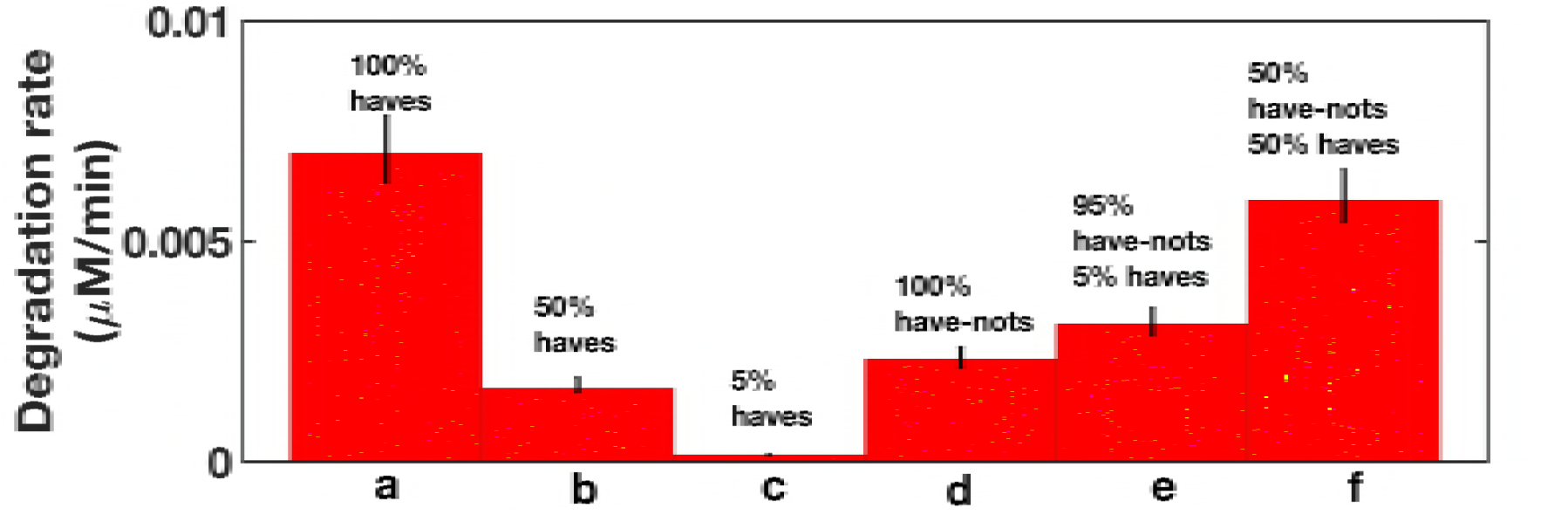
Comparison of the degradation rate of the extra cellular PDE produce by different cell populations. 100% corresponds to a cell density of about 2 × 10^5^ cells/cm^2^. Experiments performed once.

Next, to observe the effect of the heat-treated supernatant, we conditioned the low density WT cells with the heat treated supernatant, starved them for about six hours and then measured the PDE activity in the supernatant. We found that this activity (figure 9 column c) was much higher than the activity of the low density WT cells in phosphate buffer (figure 9 column b). As a check, we verified that our heat treatment denatures PDE, column a in figure 9. Finally, the degradation rate was even higher in low density populations conditioned by the calcium enriched buffer (figure 9 column d) than in just phosphate buffer. This suggested that the addition of calcium, which keeps the cells viable, induces them to produce more PDE.

**FIG. 9.**
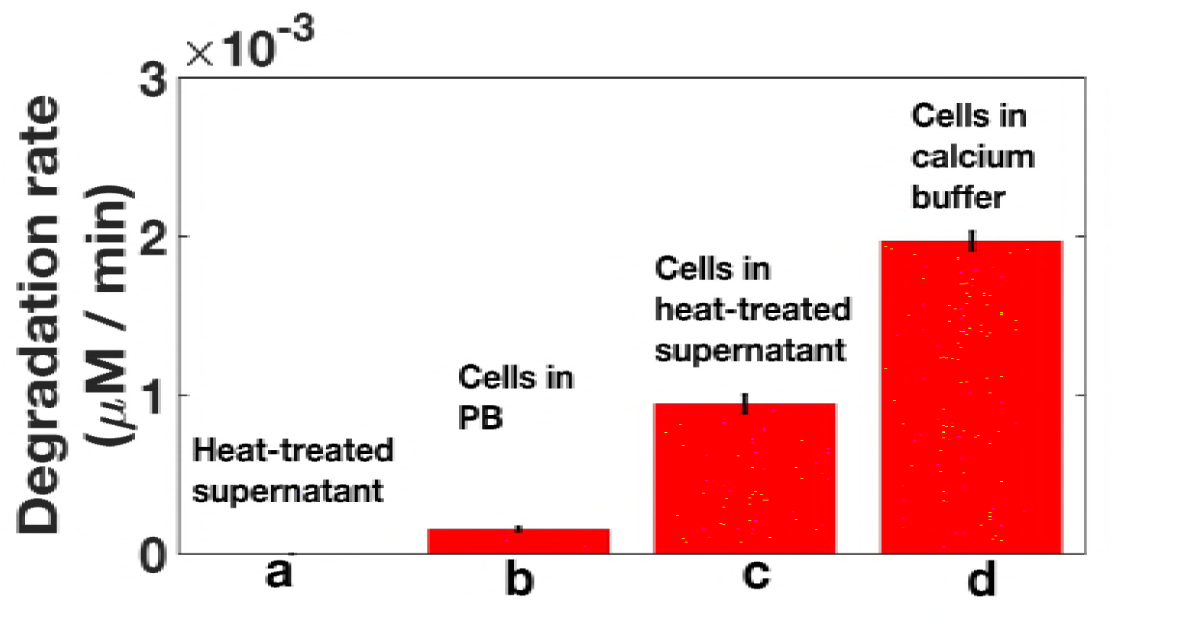
Comparison of the degradation rate of the extra cellular PDE produced by the low density populations in different conditions. PB is phosphate buffer. Low density *haves* are used in columns b, c, and d. Experiments performed once.

Our measurement of the reaction rates of the high density of *haves* is consistent with measurements by other groups [29]. However, both these rates correspond to a degradation time of about 100 min. Theoretically, wave propagation is not possible at these rates. But we do observe waves. So there must be additional sources of degradation of cAMP that these measurements missed. What these could be, is an open question.

## APPENDIX

### A1: The importance of PDE in long range signaling

Consider the typical wild type cell density used in experiments i.e. cell density *ρ* is 6 × 10^5^ cells/ml^2^. The typical cell separation *d_sep_* is given by

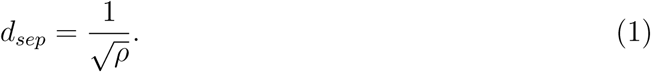

For the typical cell density mentioned above, this gives a *d_sep_* of about 13*μM*. The typical wavelength of the cAMP waves is about 1 - 2 mm. The dynamics of *D.d.* forms a reaction - diffusion system. If we assume that the degradation follows first order kinetics, the length scale *L* for diffusion is set by the diffusion constant D and the degradation rate of cAMP γ.

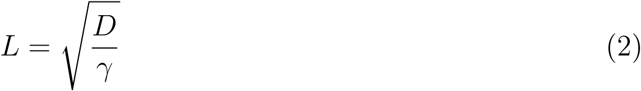

This gives,

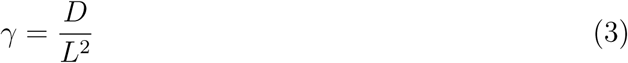

If we assume a diffusive length of about 130 *μm*, which corresponds to the distance between 10 cells (10 × d_sep_), and use D = 0.024 mm^2^/min, we get γ = 1.4 min^-1^, which corresponds to a degradation time of 0.7 min. The diffusion length scale L cannot be shorter than the typical separation between two cell, *d_sep_,* because if it were, the signal would diffuse before it reached the neighboring cell, and there would be no excitation waves. Similarly, the characteristic decay time given by 1/γ has to be shorter than the period of the waves (6 - 8 min), so that cAMP can be degraded before the next wave begins. Therefore, the value of the degradation rate at high cell densities allows wave propagation because it is shorter than the period of the wave.

Now, the density of our low density population is about 70 times smaller than the value considered above. If we assume that the degradation scales linearly with density, the degradation rate of the low density population will be 70 times smaller, and the degradation time will be 70 times larger, or about 49 min. This number is much higher than the period of the waves. This larger degradation time will not allow wave propagation. As we have seen in figure 8, the extra cellular PDE decreases more rapidly with density than a linear fit would predict. Therefore, for aggregation, more PDE is necessary in the low density *haves.*

### A2: Activity of exogenous PDE

To illustrate the calculation of activity, let us start with the activity stated on the vial of PDE. For example, in one experiment, the activity of PDE4A was 2429 units/mg. 1 unit = 1 nmol/min at 37°C. Since our experiments are at 22°C, we used 1 unit = 0.5 nmol/min.

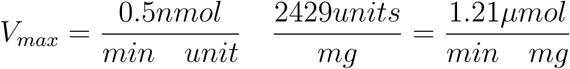

Let us assume that we dilute the sample to 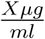. Then,

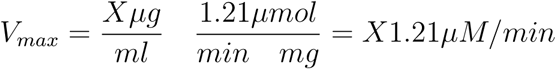

We know that the Michaelis-Menten constant for PDE4A is about *K_M_* = *5μM* [32]. At high substrate concentration, the degradation rate is given by 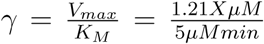. The degradation time is given by 1/γ. As a particular case, when X = 0.0032, the degradation time is 129 min, whereas when X = 20, the degradation time is 0.2 min. We don’t expect tight aggregates in the former case, whereas we do expect them in the latter case, as is shown in figure 6c.

### A3: Obtaining the degradation rate

We modeled the system in the following way. AMP is produced through the action of PDE, and there is a decay rate. For PDE produced by *D.d.*, it is known that the value of the Michaelis-Menten constant is 0.75*μ*M [29]. In our experiments, we add a much higher concentration of cAMP. So the rate of the reaction can be assumed to be independent of the substrate concentration. This rate is also *V_max_,* the maximum rate of the reaction. We observe that AMP concentration decays after the reaction is stopped by adding the stop solution. Accounting for the production and decay of AMP, we can write

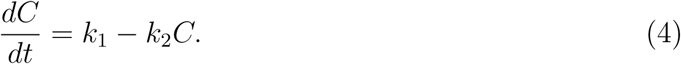

**FIG. 10.**
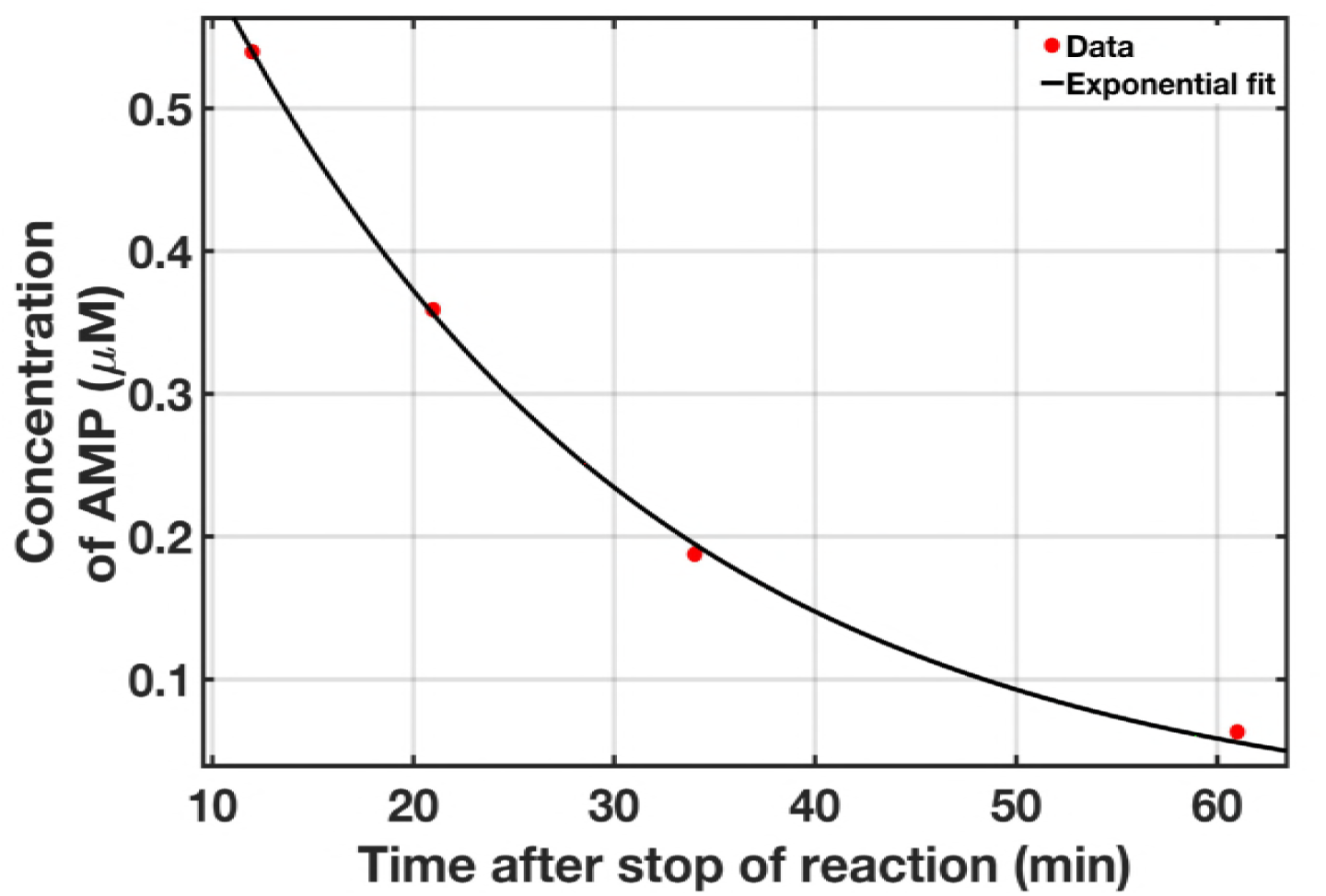
Sample fit of the decay of the concentration of AMP as a function of time.

Here C is the concentration of AMP, *k*_1_ is the degradation rate of PDE which creates AMP (this is the rate we are interested in finding) and *k*_2_ is the decay rate of AMP. The solution to the equation is

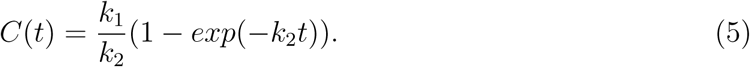

We stop the reaction after 20 min, the concentration of AMP after 20 min is C(20).

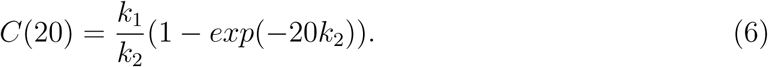

After these 20 min, there is only decay of AMP because PDE doesn’t act anymore. The equation for the concentration of AMP after these 20 min is therefore the decaying exponential

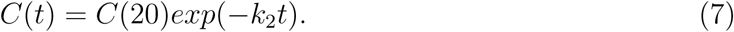

Here t is time after stopping the PDE action. We measure C(t) at different times after stopping. We first subtracted the background AMP and then converted the luminosities to concentrations using the AMP standards. Then we fit an exponential to the decaying concentration of AMP to get *C*(20) and *k*_2_.We plug these values back into equation 6 to get the decay rate *k*_1_. The error in *k*_1_ is calculated by propagating the errors in *C*(20) and *k*_2_ from the confidence intervals provided by the fit. A sample fit is shown in the figure 10.

